# Preliminary evidence of a relationship between sleep spindles and treatment response in epileptic encephalopathy

**DOI:** 10.1101/2023.04.22.537937

**Authors:** John R. McLaren, Yancheng Luo, Hunki Kwon, Wen Shi, Mark A. Kramer, Catherine J. Chu

**Affiliations:** Department of Neurology, Massachusetts General Hospital, Boston, MA, USA 02114; Harvard Medical School, Boston, MA, USA 02115; Department of Mathematics and Statistics & Center for Systems Neuroscience, Boston University, Boston, MA, USA 02215

## Abstract

**Objective:** Epileptic encephalopathy with spike wave activation in sleep (EE-SWAS) is a challenging neurodevelopmental disease characterized by abundant epileptiform spikes during non-rapid eye movement (NREM) sleep accompanied by cognitive dysfunction. The mechanism of cognitive dysfunction is unknown, but treatment with high-dose diazepam may improve symptoms. Spike rate does not predict treatment response, but spikes may disrupt sleep spindles. We hypothesized that in patients with EE-SWAS: 1) spikes and spindles would be anticorrelated, 2) high-dose diazepam would increase spindles and decrease spikes, and 3) spindle response would be greater in those with cognitive improvement.

**Methods:** Consecutive EE-SWAS patients treated with high-dose diazepam that met criteria were included. Using a validated automated spindle detector, spindle rate, duration, and percentage were computed in pre- and post-treatment NREM sleep. Spikes were quantified using a validated automated spike detector. Cognitive response was determined from chart review.

**Results:** Spindle rate was anticorrelated with spike rate in the channel with the maximal spike rate (*p*=0.002) and averaged across all channels (*p*=0.0005). Spindle rate, duration, and percentage each increased, and spike rate decreased, after high-dose diazepam treatment (*p≤*2e-5, all tests). Spindle rate, duration, and percentage (*p*≤0.004, all tests) were increased in patients with cognitive improvement after treatment, but not those without. Changes in spike rate did not distinguish between groups.

**Interpretation:** These findings confirm thalamocortical disruption in EE-SWAS, identify a mechanism through which benzodiazepines may support cognitive recovery, and introduce sleep spindles as a promising mechanistic biomarker to detect treatment response in severe epileptic encephalopathies.

## Introduction

Epileptic encephalopathy with spike wave activation in sleep (EE-SWAS) is a rare electroclinical syndrome caused by a variety of etiologies and characterized by cognitive deceleration, plateau, or regression concordant with abundant spike-and-wave pattern on EEG during non-rapid eye movement (NREM) sleep.^1-4^ The cognitive patterns can vary from mild, domain-specific deficits, to global developmental plateaus, to profound regression.^5-7^ The electrographic spike and wave activity can range from focal to generalized, and from abundant to continuous, and by definition, the epileptiform activity is potentiated during slow wave sleep compared to wakefulness. While epilepsy in these syndromes may resolve over time, the cognitive sequelae of the underlying pathophysiology are often long-lasting.^2^ The cause of cognitive dysfunction in EE-SWAS is unknown and no reliable EEG biomarkers are available to detect and track cognitive risk or treatment response.

Previous studies have shown that epileptiform activity does not reliably predict cognitive function on response to treatment in EE-SWAS.^8-10^ Several lines of research have suggested that epileptiform spikes can “hijack” the thalamocortical circuits necessary for sleep spindle production.^11, 12^ Sleep spindles are discrete 9-15 Hz oscillations generated in the thalamic reticular nucleus (TRN) during NREM sleep that support sleep-dependent memory consolidation and positively correlate with IQ, procedural memory, and semantic memory tasks in both typically developing children and those with neurodevelopmental disorders.^13-20^ Pharmacologic manipulations can transform spindles into spike and wave activity in slice preparations,^21^ computational models,^22^ and *in vivo* animal models.^23^ Consistent with these observations, children with a milder version of EE-SWAS, self-limited epilepsy with centrotemporal spikes (SeLECTS), have focal spindle deficits that anticorrelate with spike rate. In this same population, spindle rate, but not spike rate strongly correlates with IQ and performance on motor planning and attention tasks.^24, 25^

High-dose intravenous benzodiazepine therapy is a common empirical treatment for severe EE-SWAS;^26-30^ however, the mechanism behind how benzodiazepines improve neurocognitive outcome is not well understood. To evaluate for thalamocortical circuit dysfunction in EE-SWAS and understand the impact of high-dose diazepam treatment, we measured epileptiform spikes and sleep spindles using automated approaches in a cohort of patients with EE-SWAS before and after high-dose diazepam treatment. Consistent with a competitively shared thalamocortical circuitry, we hypothesized that spindle rate would be inversely related to spike rate. We further hypothesized that high-dose diazepam treatment would increase spindles and decrease spikes in patients with EE-SWAS. Lastly, we hypothesized that patients with cognitive improvement would have a larger change in spindles after high-dose diazepam. These findings would confirm thalamocortical disruption in EE-SWAS, suggest a mechanism through which benzodiazepines may support cognitive recovery, and highlight sleep spindles as a potential mechanistic biomarker to detect epileptic encephalopathy and assess treatment response.

## Methods

### Subject inclusion

A retrospective chart review was conducted for all patients with overnight EEG recordings in the Massachusetts General Hospital *for* Children Epilepsy Monitoring Unit (EMU) between 10/2009 and 2/2022. Patients were eligible for inclusion if they met the following criteria: 1) age <18 years, 2) electroclinical documentation of EE-SWAS, 3) treatment with high-dose diazepam (defined as ≥1mg/kg) for at least one night during the EMU admission, and 4) pre- and post-diazepam raw EEG data that included NREM sleep. We note that we included both NREM stages 2 and 3 together due to challenges separating the two in the presence of near-continuous spike and wave activity. For each patient, clinical information was obtained from the electronic medical record including age, sex, epilepsy diagnosis, suspected etiology, neurobehavioral comorbidities, anti-seizure medications administered during the pertinent EEG recordings, and side effects. For those who continued diazepam treatment after hospital discharge, cognitive response following diazepam treatment was obtained from chart documentation by their treating neurologist. For inclusion in the analysis of cognitive response, a minimum of four weeks of observation on high-dose valium was used. Patients who were not continued on medication after their inpatient evaluation or who were lost to follow-up before 4 weeks of observation were excluded from the analysis of cognitive response..

### EEG recordings

Clinical EEG data were acquired following the international 10-20 system for electrode placement (Fp1, Fp2, F3, F4, C3, C4, P3, P4, F7, F8, T3, T4, T5, T6, O1, O2, Fz, Cz and Pz) with a standard clinical recording system (Xltek, a subsidiary of Natus Medical). The sampling frequency was either 256 or 512 Hz. All EEG electrodes were placed by registered EEG technicians and impedances were below 10 kOhms. All available NREM 2 and 3 sleep epochs were manually clipped for the analysis according to standard criteria on the baseline night and after high-dose diazepam administration.^31^ Channels contaminated by continuous artifact were ignored and epochs with large movement artifacts were removed from the analysis. The sleep epochs were re-referenced to an average reference for subsequent analysis. All data analyses were conducted in accordance with protocols approved and monitored by the local Institutional Review Board according to National Institutes of Health guidelines.

### Manual spindle markings

Given the abundance of epileptiform spikes in our dataset, and the challenges of automatically detecting spindles in the setting of such sharp events, we manually marked spindles in a subset of our patient recordings to both further train our automated spindle detector and test its performance in these data. 1000 seconds of pre- and post-treatment EEG recordings in four EE-SWAS subjects (including four pre-treatment, four post one night of treatment, and two post two nights of treatment) were reviewed from 18 channels within the standard 10-20 EEG montage, and sleep spindles were manually marked by consensus between two reviewers (JM, YL) following standard criteria.^32^

These manual spindle markings were then used to update our automated spindle detector, validated for use in EEGs with epileptiform spikes.^24^ To do so, we computed theta power, sigma power, and the Fano factor for each 0.5 s interval (0.1 s steps) for each channel in the marked data and estimated a transition matrix between spindle and non-spindle states. These values were then used to estimate the probability of a spindle automatically (in place of manual markings). For further details on the spindle detector see Kramer et al, 2021.

### Automated spindle detection

To test the performance of the automated spindle detector, and to ensure the detector was not overfit to the EE-SWAS training data, we performed a leave-one-out cross-validation procedure, in which the original detector was updated to include training data from nine EE-SWAS recordings and then applied to the remaining tenth EE-SWAS recording. The automated detections were then compared to the manual detections for the tenth (left-out) recording. This process was repeated for all ten recordings and performance was assessed using the F1 statistic (the harmonic mean of sensitivity and specificity, a commonly applied measure of spindle detector performance).^24, 33^ This procedure was repeated to perform leave-one-out cross-validation by subject (i.e., the detector was trained on all recordings except those from an excluded subject and then tested on the recordings from the excluded subject). The automated detector trained on the entire training dataset was applied to the full experimental dataset.

### Automated spike detection

The Persyst13 (Persyst Development Corporation, San Diego) spike detector was applied to the same epochs as above to compute the spike rate per channel and averaged across all channels. This software has been previously found to be non-inferior to human experts in calculating spike wave index in patients with abundant spike activity due to EE-SWAS.^34^

### Power spectra calculations

To compare pre- and post-diazepam power spectra, we computed the power spectra using a two second sliding window (Hamming tapered) with 50% overlap in pre- and post-diazepam NREM sleep EEGs and artifact-free awake EEGs manually collected from the same subjects. The spectra in each window were normalized by the summation of power from 1 to 40 Hz and then averaged across all electrodes and all 2s windows. To evaluate for discrete changes in sigma activity during sleep (reflecting sleep spindle activity) separate from diffuse changes in beta activity observed in both wake and sleep, the power spectra from wake were then subtracted from the power spectra during sleep for each subject.

### Statistical tests

To test the relationship between spike rate and spindle rate, we modeled spindle measures (rate, duration, and percentage (*e*.*g*., the percentage of time during N2 with sleep spindles)) as a function of spike rate using the following linear mixed effects model:

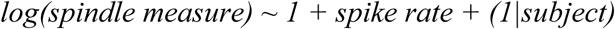

where each spindle measure was scaled to the baseline values by dividing by the measure on the pre-treatment night and (1|subject) accounts for the repeated measurements (pre- and post-treatment day 1 and day 2) for subjects.

To test the impact of high dose diazepam on spindle measures, we used the following linear mixed effects model:

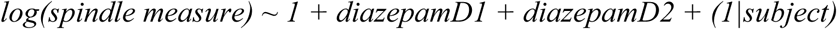

where measure is either spindle rate, spindle duration, spindle percentage (each scaled to baseline values), and diazepamD1 and diazepamD2 indicate the first and second night of treatment, respectively.

To test the impact of high dose diazepam on spike rate, we used the following linear mixed effects model:

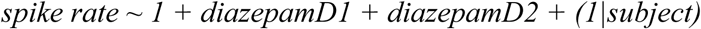

where spike rate was scaled to baseline values, and diazepamD1 and diazepamD2 indicate the first and second night of treatment, respectively.

To test for a difference in response of spindles and spikes after high-dose diazepam between those with and without cognitive improvement, we used Wilcoxon signed rank tests of the measure scaled to the rates on the baseline night and then compared to 0 for spindle features which were first log-transformed (right-tailed) or 1 (spike rate, left-tailed) (*e*.*g*., no change). To evaluate for a larger change in values in children with cognitive improvement compared to no improvement, we used a left-tailed Ranksum test.

### Data availability

Raw data were generated at Massachusetts General Hospital and the Athinoula A. Martinos Center for Biomedical Imaging. Derived data supporting the findings of this study are available from the corresponding author on request. Software for the detection of spindle events is available for reuse and further development at https://github.com/Mark-Kramer/Spindle-Detector-Method.

## Results

### Clinical and demographic characteristics

Data were included from forty overnight EEG recordings in seventeen patients (seventeen baseline recordings, seventeen post diazepam day one recordings, six post-diazepam day two recordings). NREM sleep EEG overnight recordings were available for analysis with mean duration 280 min (range 18-612 min). We note that NREM duration did not predict spindle rate at the channel with the maximal spike rate or averaged across all channels (p>0.5, linear models). Pretreatment cognitive comorbidities were present in all subjects, of whom formal pre- and/or post-treatment neuropsychological evaluations were available in thirteen (76%) subjects. Diazepam was not continued in three subjects after discharge and one subject was lost to follow up after 3 weeks. Post-diazepam cognitive improvements were documented by the treating neurologist in nine of thirteen subjects (69%) who were continued on treatment with diazepam after discharge. Post-diazepam transient or persistent side effects were documented in eleven (65%) subjects. Subject characteristics are listed in Table 1.

### Spindle detector validation

A total of 10,168 sleep spindles were manually marked for detector training and validation, including 3,835 spindles from four pre-diazepam EEG recordings and 6,333 spindles from six post-diazepam EEG recordings. Using leave-one-out cross-validation, the automated spindle detector had excellent performance against hand-markings across subjects and nights (F1=0.66 across EEGs and F1=0.66 across subjects) using a 95% threshold. To confirm that the detector performed similarly on data before and after high-dose diazepam, we analyzed the sensitivity and false positive rate of the detector for the baseline nights and the post-diazepam nights separately. We found no evidence of a difference in performance across the two conditions (sensitivity, *p*=0.7; FPR, *p*=0.8, two-tailed t-tests). We note that this performance is higher than prior reported automated spindle detectors.^24, 33^ The probability threshold of 95% was used for all subsequent analyses. Example detections and detector performance are shown in Figure 1.

**Figure 1.**
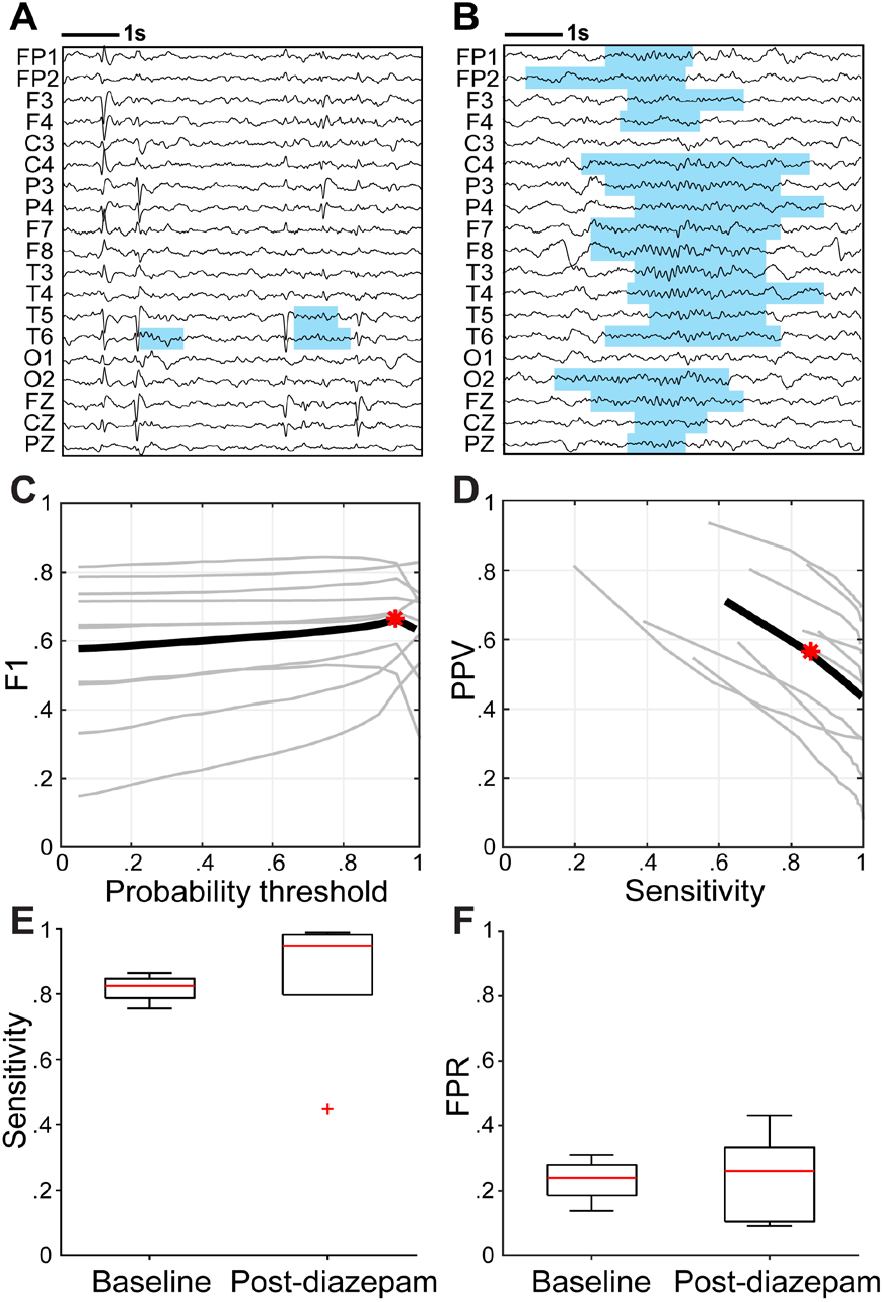
Spindle detector example and performance. (A)Example spindle detections in a baseline and (B) post-diazepam sleep EEG recording show increased spindles and decreased spikes diffusely after high-dose diazepam treatment. (C-D) Automated sleep spindle detector performance using leave-one-out procedure against manual markings in ten EEG recordings. (C) Using leave-one out cross validation, the optimal F1 statistic against hand markings was achieved at the 95% probability threshold (0.66). Grey curves represent detector performance for individual EEGs across probability thresholds and the black curve indicates the mean performance across EEGs. (D) The sensitivity and positive predictive value (PPV) of the detector at different probability thresholds (95% threshold indicated in red). (E) We find no difference in the point-wise sensitivity or (F) false positive rate (FPR) in spindle detections between EEGs from baseline nights (N=4) and post-diazepam nights (N=6).

### Spike rate and spindle activity are anticorrelated

Spike rate and spindle activity (scaled to baseline night for each patient) in the channel with the maximal spike rate and averaged across all channels were evaluated using linear mixed-effect models (Figure 2). In the channel with the maximal spike rate, log(spike rate) was anticorrelated to spindle rate (*p*=0.0019, *effect size*=-1.27, 95% CI [-2.04, -0.50]), duration (*p*=5e-6, *effect size*=-0.84, 95% CI [-1.15, -0.52), and percentage (*p*=7e-5, *effect size*=-2.10, 95% CI [-3.04, -1.15]). Averaged across all channels, log(spike rate) was anticorrelated to spindle rate (*p*=5e-4, *effect size*=-1.48, 95% CI [-2.25, -0.70]), duration (*p=*5e-8, *effect size*=-0.99, 95% CI [-1.29, -0.70]), and percentage (*p*=4e-6, *effect size*=-2.38, 95% CI [-3.28, -1.49]). Thus, spike rate and spindle activity (rate, duration, and percentage) were anticorrelated in patients with EE-SWAS.

**Figure 2.**
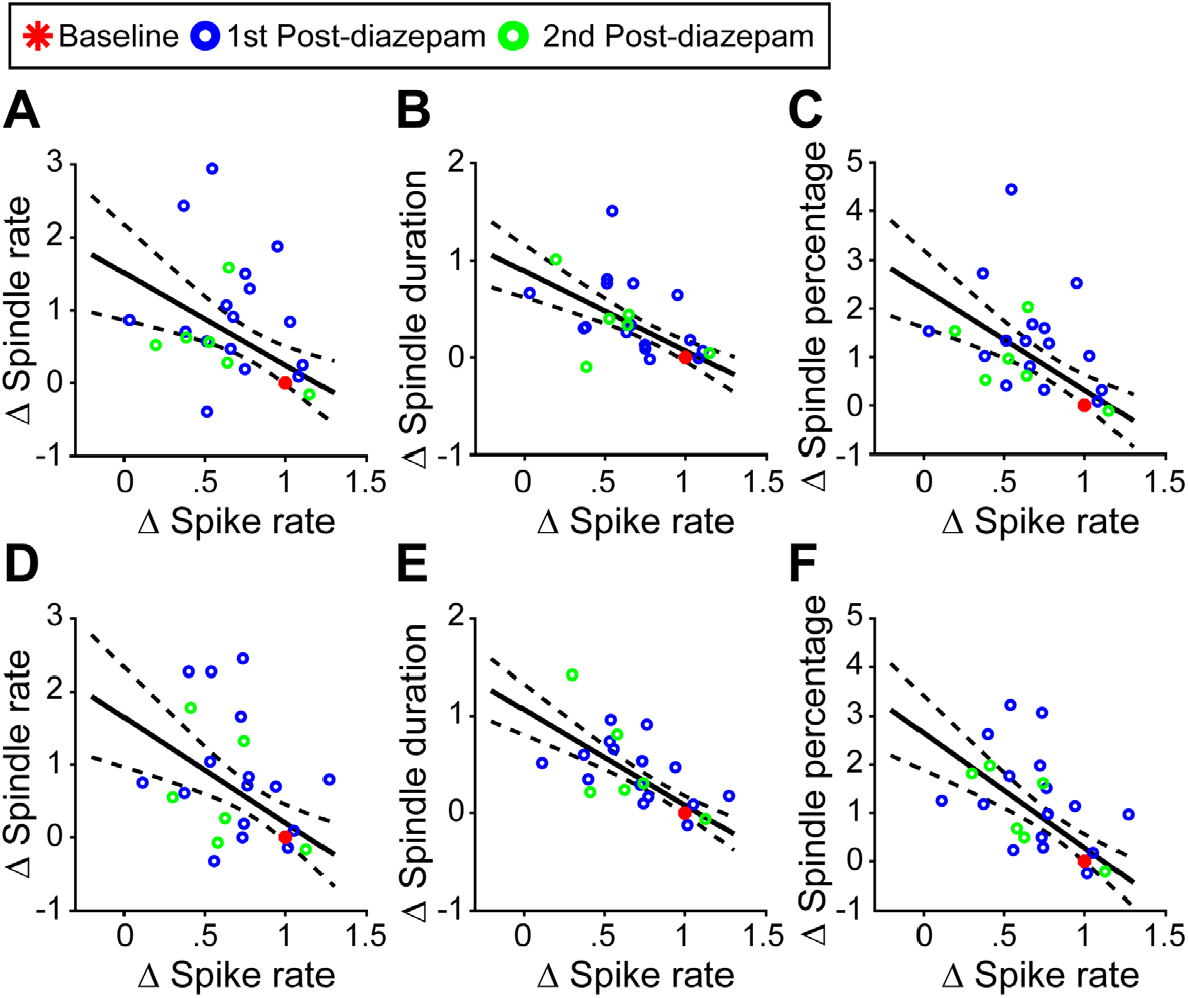
Spindles and spikes are anticorrelated. Each point represents an individual night of EEG recording. (A) log(spindle rate), (B) log(spindle duration), and (C) log (spindle percentage) are each anticorrelated with spike rate in the channel with the maximal spiking rate (both spindle and spike values are scaled to baseline rates on the pre-diazepam treatment night). (D-F) Similar plots showing average rates across all EEG channels. All relationships, p<0.001.

### High-dose diazepam increases sleep spindles

Post-diazepam spindle metrics (scaled to pre-diazepam baseline values for each patient) in the channel with the maximal spike rate and averaged across all channels were evaluated using a linear mixed-effect model (Figure 3). In the channel with the maximal spike rate, spindle rate (*p*=3e-6, *effect size*=1.02 95% CI [0.64, 1.40), duration (*p*=1e-5, *effect size*=0.43, 95% CI [0.25, 0.60]), and percentage (*p*=4e-7, *effect size*=1.45, 95% CI [0.97, 1.93]) each increased on the first night after diazepam treatment compared to baseline. In the channel with the maximal spike rate, spindle rate (*p*=0.026, *effect size*=0.62, 95% CI [0.08, 1.16]), duration (*p*=0.003, *effect size*=0.39, 95% CI [0.14, 0.64]), and percentage (*p*=0.005, *effect size*=0.99, 95% CI [0.31, 1.67]) each further increased after a second night of high-dose diazepam treatment compared to baseline. Averaged across all channels, we found similar results: spindle rate (*p*=2e-5, *effect size*=0.88, 95% CI [0.52, 1.24]), duration (*p*=1e-6, *effect size*=0.44, 95% CI [0.29, 0.60]), and percentage (*p*=3e-7, *effect size*=1.31, 95% CI [0.89, 1.74]) increased on the first night after diazepam treatment. Following the second night of high-dose diazepam treatment, average spindle rate (*p*=0.01, *effect size*=0.66, 95% CI [0.14, 1,18]), duration (*p*=5e-5, *effect size*=0.51, 95% CI [0.28, 0.74]), and percentage (*p*=7e-4, *effect size*=1.12, 95% CI [0.51, 1.74]) further increased.

**Figure 3.**
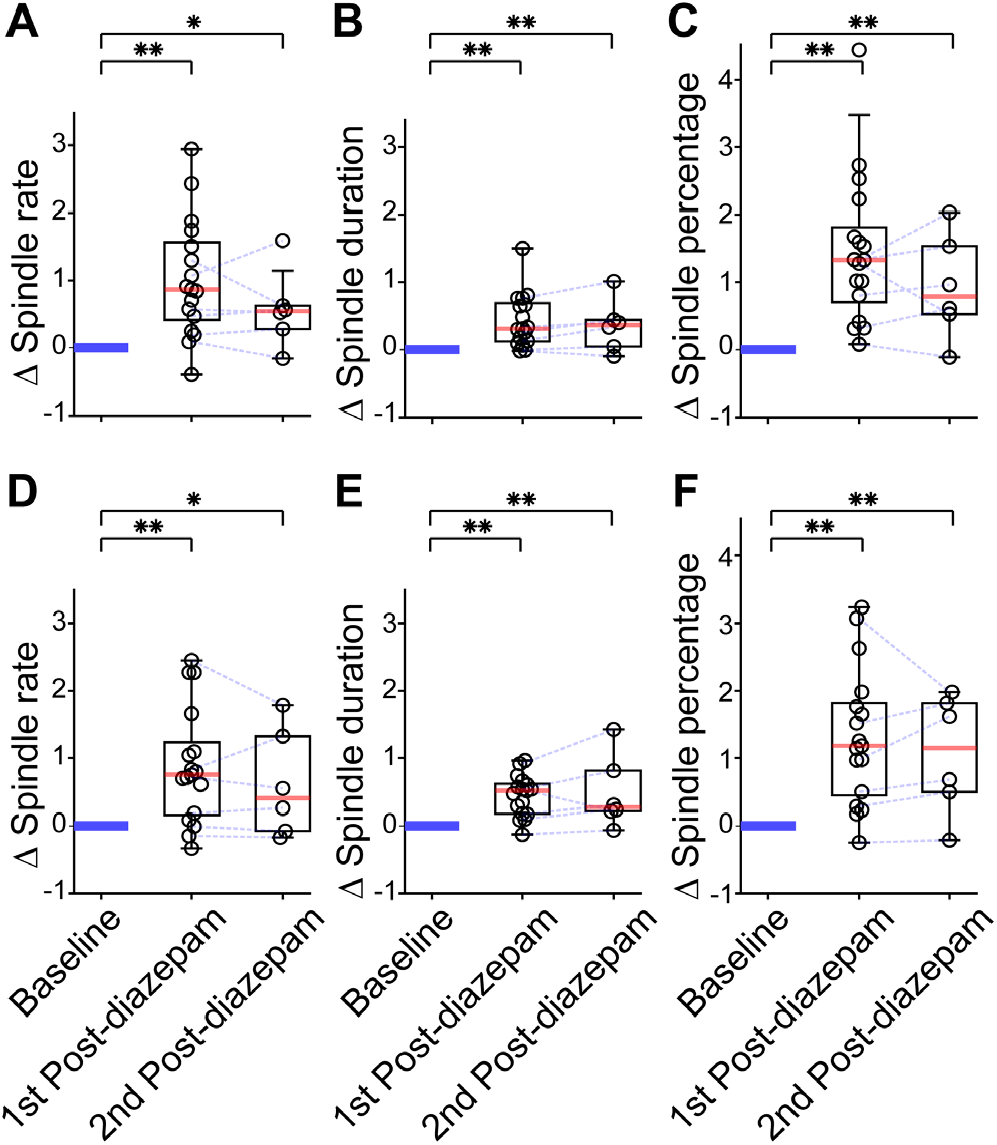
Spindle rate, duration, and percentage are increased after high-dose diazepam. In the channel with the maximal spiking rate, (A) log(spindle rate), (B) log(spindle duration), and (C) log (spindle percentage) increase after one night of high-dose diazepam and further increase in subjects who received a second night of treatment (all values are scaled to the baseline values; blue lines connect value across nights from the same subject). (D-F) Consistent findings are observed averaged across all channels. *p<0.05, **p<0.001.

As benzodiazepines are known to cause a generalized increase in beta activity (12.5-30 Hz) in the EEG,^35^ we subtracted the spectra in wake from sleep following high-dose diazepam treatment to confirm a specific increase in the pediatric spindle frequency bands (9-15 Hz) during sleep. To do so, we analyzed equivalent durations of sleep and awake recordings from twelve subjects with available artifact-free wake EEGs (total 300 min NREM and 300 min wake). Following high-dose diazepam treatment, there was a discrete increase in 11-14 Hz power during NREM sleep that overlapped with the sigma bump present during NREM sleep in the baseline night (Figure 4). Taken together with the results of our spindle detector, these data provide additional evidence for an increase in sleep spindle activity after high-dose diazepam treatment.

**Figure 4.**
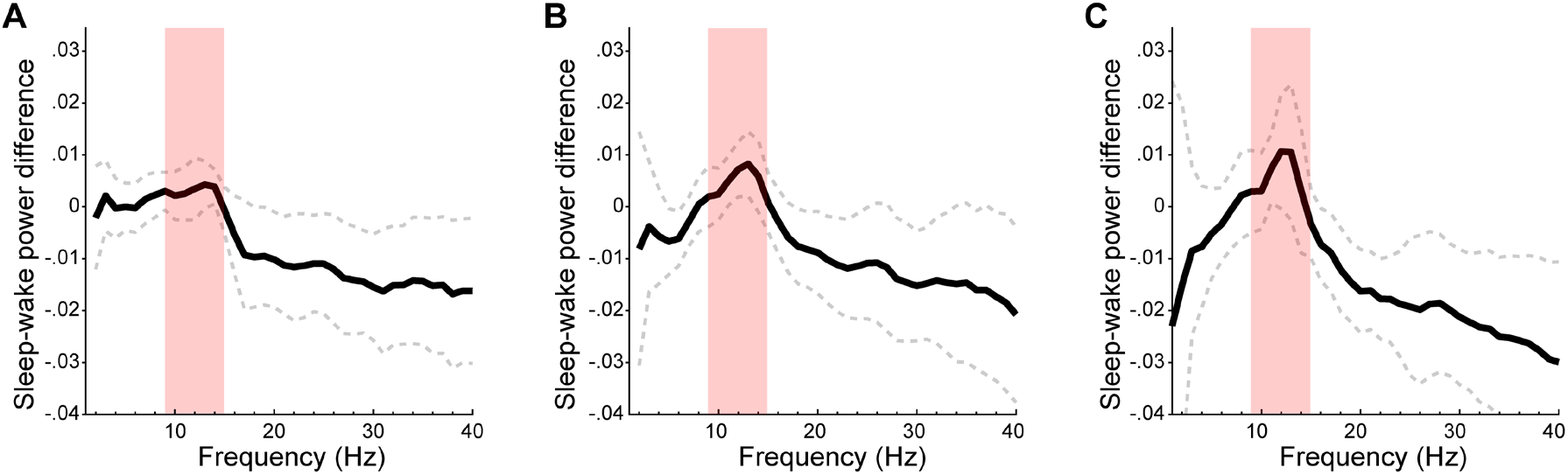
Increase in sigma band power during N2 sleep following diazepam treatment. The mean difference in power between sleep and wake is plotted across frequencies on left) the baseline night middle) after on the first night after high dose diazepam treatment, and right) on the night after a second dose of high dose diazepam treatment. A discrete increase in sigma band power (red shading) is seen after each night of high-dose diazepam treatment, corresponding to increases in sleep spindles. Solid (dashed) curves indicate mean (95% CI) across subjects.

### High-dose diazepam decreases spike rate

Pre- and post-diazepam spike rates (scaled to baseline for each patient) in the channel with the maximal spike rate and averaged across all channels were evaluated using a linear mixed-effect model. In the channel with the maximal spike rate, spike rate decreased on the first night after diazepam treatment compared to baseline (*p*=1e-4, *effect size*=-0.33, 95% CI [-0.48, -0.17]) and remained lower or further decreased after a second night of high-dose diazepam treatment (*p*=3e-4, *effect size*=-0.41, 95% CI [-0.62, -0.20]). Averaged across all channels, spike rate decreased following the first night of high dose diazepam treatment (*p*=3e-4, *effect size*=-0.30, 95% CI [-0.45, -0.15]) and remained lower or further decreased following a second night of high dose diazepam treatment (*p*=8e-4, *effect size*=-0.37, 95% CI [-0.57, -0.17]) (Figure 5). Taken together, these results suggest high-dose diazepam decreases spike rate focally and globally in children with EE-SWAS from a variety of etiologies.

**Figure 5.**
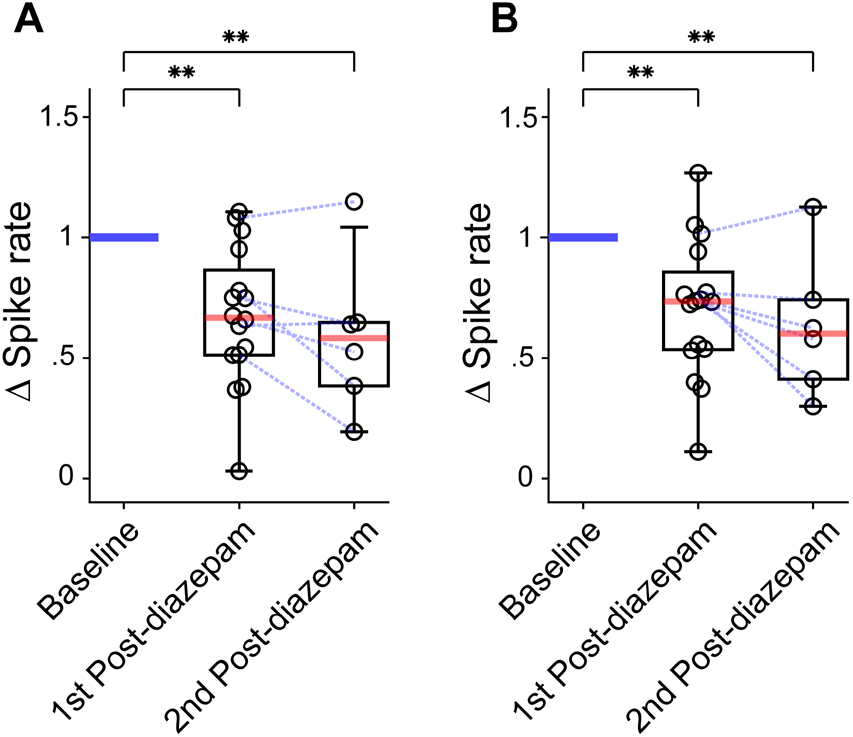
Spike rate is decreased after high-dose diazepam. (A) In the channel with the maximal spiking rate, spike rate is decreased after one night of high-dose diazepam and further decreased after a second night. All values are scaled to the baseline values; blue lines connect value across nights from the same subject). (B) Similar results are seen averaged across all EEG channels. *p<0.05, **p<0.001.

### Sleep spindles are increased in patients with cognitive improvement after high-dose diazepam

We evaluated whether children with (*n*=9) or without (*n*=4) cognitive improvement after treatment had increases in spindle metrics compared to baseline (Figure 6). Children who were not continued on high-dose benzodiazepine after discharge (n=3) or lost to follow-up (1 who moved out of the country 3 weeks after discharge) were not included in this analysis. We found that the children with cognitive improvement had an increase in spindle rate, duration, and percentage after diazepam treatment in both the maximal spiking channel (*p*=0.004, *p*=0.004, *p*=0.002) and averaged across all channels (*p*=0.004, *p*=0.002, *p*=0.004). In contrast, children with no improvement after diazepam had no change in spindle rate, duration, or percentage in the channel with the maximal spike rate or averaged across all channels (*p*>0.12, all tests). Compared to children without cognitive improvement, those with cognitive improvement had a larger change in spindle rate in the maximal spiking channel and averaged across all channels (*p*=0.017, *p*=0.025) and a larger change in spindle percentage in the maximal spiking channel (*p*=0.017). Thus, children with cognitive improvement after high-dose diazepam, but not those without, had an increase in spindle rate, duration, and percentage from baseline after treatment. Further, the change in spindle rate and percentage after treatment were significantly higher in children with cognitive improvement.

**Figure 6.**
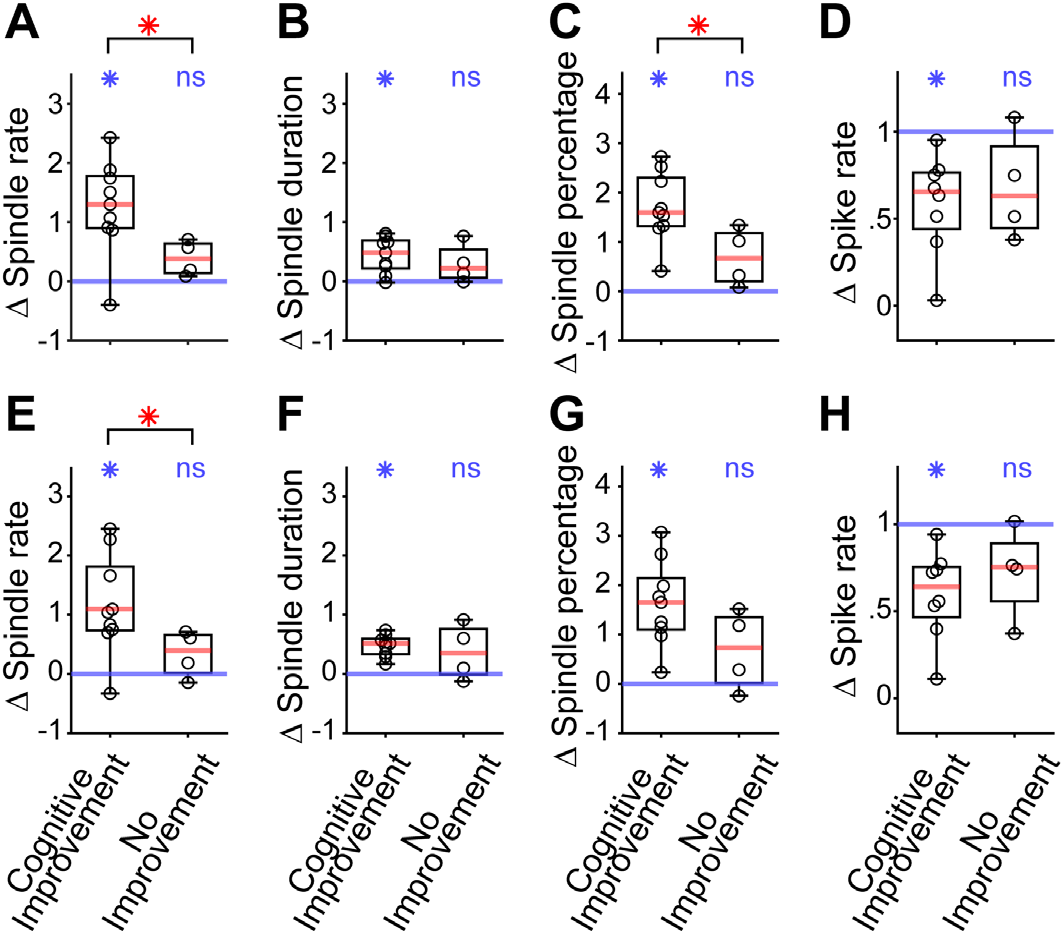
Post-diazepam sleep spindle response, not spike response, correlates with cognitive improvement. In the channel with the maximal spiking rate, (A) spindle rate, (B) duration, and (C) percentage are each increased from baseline and (D) spike rate is decreased from baseline in children with cognitive improvement. Larger changes in (A) spindle rate and (C) percentage after high-dose diazepam treatment are observed in children with cognitive improvement than children without cognitive improvement. (E-H) Similar results from baseline are seen averaged across all EEG channels. Only changes in spindle rate are different between groups. *p<0.05.

### Spikes are reduced in patients with cognitive improvement

We evaluated whether children with or without cognitive improvement after high-dose diazepam treatment had decreases in spike rate compared to baseline. We found that children with, and not those without, cognitive improvement had decreases in spike rate in the channel with the maximal spike rate (*p*=0.004, *p*=0.125) and averaged across all channels (*p*=0.004, *p*=0.125) compared to baseline. No difference in the degree of change in spike rate between the two groups was detected (p>0.6).

## Discussion

Epileptic encephalopathy with spike wave activation in sleep is a challenging condition with heterogeneous etiologies that features cognitive plateau, deceleration, or regression concomitant with the emergence of abundant epileptiform activity during NREM sleep.^1-4^ Despite this diagnostic association, spike rate does not reliably predict cognitive symptoms and a joint pathophysiological mechanism leading to both the ongoing seizures and cognitive deficits remain unknown. Using validated and reproducible methods, we found that, in a cohort of patients with severe EE-SWAS from a variety of etiologies, spike rate and spindle activity were anticorrelated. In addition, for this same cohort, we found that high-dose diazepam decreased spike rate and increased spindle rate. Finally, we found that patients with cognitive improvement after treatment had larger changes in spindle metrics than spike rates, compared to those without cognitive improvement. Taken together, these data support increased sleep spindles as a mechanism underlying cognitive improvement after high-dose diazepam treatment in EE-SWAS and introduce spindles as a cognitive biomarker to help diagnose and detect treatment response in this challenging disease.

In our study, we updated an automated spindle detector previously designed to reduce spurious spindle detections in the setting of epileptiform spikes.^24^ After retraining this detector using an additional >10,000 spindles marked in EE-SWAS datasets, we found that the detector had excellent performance in a heterogeneous EE-SWAS patient population. The use of an automated strategy for spindle detection eliminates subjectivity in measurements and enables consistent and reproducible measures of sleep spindles even in this challenging dataset.

The thalamocortical circuit underlying sleep spindles has been well characterized. Spindles are generated in the thalamic reticular nucleus (TRN) which is comprised entirely of GABAergic neurons.^36-40^ These neurons project primarily to glutamatergic thalamocortical neurons, which entrain cortical areas to their sigma-frequency rhythms.^11^ The resulting rhythms - sleep spindles - are amplified and propagated through corticocortical and thalamocortical circuits. Corticothalamic neurons send glutamatergic inputs back to the thalamus, producing a feedback loop regulated primarily by GABAergic and glutamatergic neurotransmission. Benzodiazepines enhance the binding of GABA to the GABAA receptor, enhancing GABAergic hyperpolarization, which in a computational model was expected to reduce epileptiform spikes and enhance spindles at the TRN.^22^ Our study empirically demonstrates that high-dose diazepam does in fact modulate this thalamocortical circuit to support spindles and reduce spikes.

Several lines of evidence suggest that thalamocortical circuits are disrupted in sleep-activated spike syndromes. In SeLECTS, focal microstructural^41^ and macrostructural^42^ lesions have been observed in Rolandic thalamocortical white matter. Similarly, thalamic lesions are more commonly found in children with epileptic encephalopathy with spike wave activation in sleep (EE-SWAS) compared to patients with developmental regression without EE-SWAS,^43^ and patients with unilateral spikes in EE-SWAS are more likely to have thalamic lesions lateralized to the affected side.^44^ The data we present here further supports the growing evidence of thalamocortical circuit disruption in patients with EE-SWAS from different etiologies (lesional, genetic, and idiopathic), impacting different brain regions, and in the setting of varying medication regimens, highlighting a similar pattern across a heterogeneous population. Specifically, we show that epileptiform spike and sleep spindle rates are anticorrelated. The involvement of sleep spindles confirms thalamocortical circuit involvement and suggests that a competitive imbalance between spikes and sleep spindles could result in both the seizures and cognitive symptoms observed in epileptic encephalopathies.

Although studies evaluating the relationship between spikes and cognitive function have reported inconsistent results,^9, 10, 27^ the role of sleep spindles in sleep-dependent learning and memory processes is well established in adults and children.^13-20^ Further supporting the mechanistic role of this rhythm in memory processes, both pharmacologic^45, 46^ and non-pharmacologic^47-50^ interventions that increase spindle activity result in improved sleep-dependent memory consolidation, while interventions that decrease spindle activity impair memory.^51^ We previously reported a single case of a patient with idiopathic EE-SWAS in whom post-diazepam increased sleep spindle rates coincided with dramatically improved scores on multiple neurocognitive tests.^29^ As observed in the larger cohort reported here, high-dose diazepam did not result in improved cognition in all subjects; one large, pooled analysis reports improvement in 68% of children.^52^ However, we found that spindle response, not spike response, was different between outcome groups. Although spikes and spindles are anticorrelated, we note that the confidence intervals for the effect size approach (but do not include) zero. We note that given the small cohort analyzed here we did not have sufficient power to detect a small effect. However, within even this small cohort, we detected a significant difference in spindles between groups. Analysis from a larger sample of subjects may help identify a weaker relationship between spike rate and cognitive function. The findings here, and those in our prior work,^24, 29^ that spindles predict cognitive response better than spikes, suggest that spikes do not directly impact cognitive function. Rather, some spikes may interfere with sleep spindles, which more directly impact cognitive function. Thus, sleep spindles may provide a reliably detected, sensitive, more direct mechanistic biomarker for cognitive symptoms and treatment response in EE-SWAS. Further, given that there are no current FDA-approved treatments for cognitive symptoms in epileptic encephalopathy, this mechanistic biomarker could provide a much-needed screening target against which to test developing therapeutics to improve cognitive symptoms in this heterogeneous disorder.

We note that due to the retrospective nature of our study, we did not have quantitative measurements pre and post high-dose diazepam treatment to examine the specific relationship between spindle responses and the degree or nature of cognitive improvement. We also recognize that the appropriate quantitative measure for cognitive response related to sleep spindles is a measure of sleep dependent memory consolidation^16, 53-55^ which are not available in routine neuropsychological assessments or standard clinical exams, but long-term assessments that include periods of sleep are more likely to capture these improvements.^56^ To categorize clinical outcome here, we relied on the long-term qualitative assessments of the treating providers, consistent with current clinical practice in this ultra-rare epilepsy syndrome^57^. That the subjects included here had clinical responses determined by six separate attending neurologists before the conception of this study increases the generalizability and rigor of our findings. As benzodiazepines can also introduce side effects of sedation and psychomotor impairment, it will be important to evaluate for improved cognitive function alongside the improvements in sleep spindles when used in treatment. This work provides the hypotheses and motivation for these questions to be answered in future prospective studies.

## Acknowledgements

This work was supported by NINDS R01NS115868

## Author Contributions

John R. McLaren: acquisition and analysis of data, drafting a significant portion of the manuscript/figures

Yancheng Luo: acquisition and analysis of data, drafting a significant portion of the manuscript/figures

Hunki Kwon: analysis of data, drafting a significant portion of the manuscript/figures

Wen Shi: analysis of data, drafting a significant portion of the manuscript/figures

Mark A. Kramer: analysis of data, drafting a significant portion of the manuscript/figures

Catherine J. Chu: conception and design of the study, analysis of data, drafting a significant portion of the manuscript/figures

